# *Drosophila* glutathione S-transferase S1 exacerbates phenotypes of the gain-of-function voltage-gated sodium channel mutant *para^Shu^* through a hemocyte-mediated mechanism

**DOI:** 10.64898/2026.09.26.754684

**Authors:** James Mrkvicka, Wayne Johnson, Junko Kasuya, Toshihiro Kitamoto

**Author notes:** Corresponding author: Toshihiro Kitamoto, Department of Anesthesia, Carver College of Medicine, University of Iowa, 1-675 Bowen Science Building, 51 Newton road, Iowa City, IA 52242. Department of Neuroscience & Pharmacology, Carver College of Medicine, University of Iowa, Iowa City, USA.

## Abstract

*Drosophila* glutathione S-transferase S1 (GstS1) is a unique member of the glutathione S-transferase family and is considered the ortholog of vertebrate hematopoietic prostaglandin D synthase based on amino acid sequence similarity. A previous forward genetic screen revealed that a null mutation in *GstS1* dominantly suppresses genetically induced behavioral hyperexcitability in several seizure-prone *Drosophila* mutants, including *para^Shu^*, a gain-of-function mutant of the voltage-gated sodium channel gene *paralytic* (*para*), and *eas^2^*, a loss-of-function mutant of the ethanolamine kinase gene *easily shocked* (*eas*). Here, using the GAL4/UAS system combined with *GstS1* RNAi, we show that hemocyte-specific *GstS1* knockdown suppresses hyperexcitable phenotypes in both *para^Shu^* and *eas^2^*. Notably, suppression of adult *para^Shu^* phenotypes required *GstS1* knockdown before adult eclosion, suggesting that GstS1 modulates adult neuronal excitability by influencing neural development through neuroimmune interactions. Consistent with this model, hemocyte-specific *GstS1* knockdown rescued abnormal dendritic morphology in developing class IV dendritic arborization (C4da) neurons of *para^Shu^* larvae. Moreover, *para^Shu^*and *eas^2^* larvae exhibited increased numbers of sessile hemocytes surrounding C4da neuron somata, a phenotype reversed by either *GstS1* knockdown or dietary supplementation with the ω-3 polyunsaturated fatty acid α-linolenic acid. Together, these findings reveal a previously unrecognized developmental role for GstS1 in promoting nervous system hyperexcitability through hemocyte-mediated neuroimmune interactions, potentially involving bioactive lipid signaling. Our results highlight neuroimmune crosstalk as an important mechanism linking genetic perturbations to altered neural development and seizure-associated hyperexcitability.

## 1. Introduction

More than 1,700 epilepsy-associated mutations have been identified in the human *SCN1A* gene, which encodes the α-subunit of the voltage-gated sodium channel (VGSC) Nav1.1 and is considered the most clinically relevant epilepsy gene (Kluckova et al., 2020). Notably, individuals carrying the same epilepsy-causing mutation often exhibit substantial variability in disease severity and clinical presentation (Klassen et al., 2011; Ng et al., 2024). This phenotypic heterogeneity suggests that factors beyond the primary pathogenic mutation—such as genetic variants in other genes and environmental factors—play critical roles in shaping disease outcomes. The fruit fly *Drosophila melanogaster* provides a powerful experimental system for systematically studying such phenotypic modifiers. Key advantages of this model include an extensive genetic toolkit, well-annotated genomic resources, quantifiable behavioral and electrophysiological assays, strong evolutionary conservation of epilepsy-relevant genes, controlled genetic backgrounds, a short life cycle, high fecundity, minimal ethical concerns, and low maintenance costs (Fischer et al., 2023).

*para^Shu^* is a dominant gain-of-function allele of *paralytic* (*para*), the sole VGSC gene in *Drosophila*, and causes pronounced neuronal and behavioral hyperexcitability (Kaas et al., 2016; Williamson, 1982). In a previous study, we made the serendipitous observation that flies reared on two commonly used standard food formulations exhibited marked differences in the severity of *para^Shu^* phenotypes (Kasuya et al., 2019). This observation ultimately led to the identification of α-linolenic acid (ALA) as a potent dietary suppressor of seizure-like hyperexcitability in *para^Shu^* and other seizure-prone *Drosophila* mutants (Kasuya et al., 2023). ALA is an ω-3 polyunsaturated fatty acid (PUFA) known to modulate innate immune and inflammatory responses in mammals, largely through its metabolic balance with ω-6 PUFAs and their downstream pro-inflammatory lipid mediators, such as prostaglandin E2 (Calder, 2017).

We previously conducted an unbiased forward genetic screen in *para^Shu^* mutants to identify genetic variants that phenocopy the suppressive effects of dietary ALA on genetically induced hyperexcitability. The screen identified a null allele of *Glutathione S-transferase S1* (*GstS1^M26^*), which dominantly suppresses the hyperexcitable phenotypes of *para^Shu^* and other seizure-prone mutants, similar to the effects of dietary ALA supplementation (Chen et al., 2020). These findings were further supported by RNA interference (RNAi) experiments showing that ubiquitous knockdown of *GstS1* using the *da-GAL4* driver reduced the severity of *para^Shu^* phenotypes to a degree comparable to that observed in *GstS1^M26^* mutants (Chen et al., 2020).

Glutathione S-transferases (GSTs) are a superfamily of phase II metabolic enzymes that catalyze the conjugation of glutathione (GSH; γ-glutamyl-cysteinyl-glycine) to a wide range of electrophilic xenobiotic and endogenous substrates. This reaction increases substrate solubility and promotes excretion, thereby serving primarily as a detoxification mechanism (Salinas & Wong, 1999; Vaish et al., 2020). In addition to this canonical role, GSTs perform diverse non-detoxification functions, including redox regulation, signal transduction, metabolite transport, and the metabolism of prostaglandins and leukotrienes (Hayes et al., 2005). Notably, insect GstS1 is considered an ortholog of vertebrate hematopoietic prostaglandin D synthase (H-PGDS), an enzyme that catalyzes the production of prostaglandin D_2 (_PGD_2)_, a lipid mediator involved in inflammation and a broad range of physiological processes (Sajjadian et al., 2020; Scarpati et al., 2019).

The present study represents an important step toward elucidating how loss of *GstS1* function suppresses the neurological phenotypes of seizure-prone mutants such as *para^Shu^*, and how *GstS1*-dependent mechanisms intersect with those underlying the effects of dietary ALA supplementation. Our findings identify *GstS1* as a genetic modifier of neuronal and behavioral hyperexcitability that act through macrophage-like hemocytes, highlighting a previously underappreciated role for peripheral immune cells in shaping neural development and excitability. These effects are likely mediated by bioactive lipid signaling pathways, providing mechanistic insight into how immune–lipid interactions modify seizure susceptibility. Together, our results support a developmental neuroimmune framework in which immune cell–derived signals act as key modifiers of genetically encoded neuronal dysfunction, offering new perspectives on how genetic and environmental factors converge through immune mechanisms to regulate hyperexcitable phenotypes.

## 2. Materials and Methods

### 2.1. Fly stocks and culture conditions

The Canton-S (CS) strain was used as the wild-type (WT) control. *para^Shu^*, originally referred to as *Shudderer* (*Shu*) (Williamson, 1982), was obtained from Mr. Rodney Williamson (Beckman Research Institute of the Hope, CA). Mutants of *easily shocked* (*eas^2^*) and *Glutathione S-transferase S1* (*GstS1^M26^*) were obtained from Dr. Chun-Fang Wu and Dr. Tina Tootle (University of Iowa, Iowa City, IA), respectively. *ppk-CD4-tdGFP* was a gift from Dr. Chun Han (Cornell University, Ithaca, NY). Other fly stocks, *elav-GAL4*, *He-GAL4*, *repo-GAL4*, *nSyb-GAL4*, *Lsp-GAL4*, *srpHemo-GAL80*, and *UAS-RedStinger*, were obtained from the Bloomington *Drosophila* Stock Center (Indiana University, Bloomington, IN).

Flies were reared at 25°C, 65% humidity in a 12 hr light/dark cycle on the diet originating from a recipe developed by Edward Lewis (Lewis, 1960) and modified by Rodney Williamson (Beckman Research Institute of the City of Hope, Duarte, CA): a cornmeal/glucose/yeast/agar medium supplemented with the mold inhibitor methyl 4-hydroxybenzoate (0.05 %), propionic acid, and phosphoric acid. The exact composition of the fly food used in this study was described in our previous paper (Kasuya et al., 2019) [7].

### 2.2. Analyses of para^Shu^ and eas^2^ phenotypes

#### 2.2.1. Scoring abnormal wing posture

Newly eclosed *para^Shu^* female heterozygotes (*para^Shu^*/+) were collected and assessed for abnormal wing posture either 24 hours or 5 days after eclosion. Abnormal wing posture was defined by non-overlapping (downturned) wings, as previously described (Kaas et al., 2016).

#### 2.2.2. Heat-induced seizure assay

Newly eclosed *para^Shu^* female heterozygotes were collected in groups of approximately 20 and aged for 3 to 5 days before undergoing the heat-induced seizure assay, as previously described (Sun et al., 2012). Briefly, a single fly was placed in a 15 x 45 mm glass vial (Thermo Fisher Scientific, Waltham, MA) and allowed to acclimate at room temperature for 2 to 10 min. Each glass vial containing a fly was then submerged in a water bath at 37°C for 2 min. During this period, the fly was video recorded and assessed every 5 seconds for seizure behavior, which was defined as loss of standing posture with leg shaking and/or wing beating.

#### 2.2.3. Bang sensitivity assay

This assay was performed according to a previously described protocol (Zhang et al., 2002). *eas* mutant males (*eas^2^*/*Y*) were collected in groups of approximately 10 within 24 hours after eclosion and aged for 2-3 days. Before testing, individual flies were transferred into clean 12 x 75 mm glass tubes (VWR, West Chester, PA) containing food and allowed to acclimate for approximately 30 minutes. The flies were then vortexed at maximum speed for 10 seconds, after which their recovery time was recorded. Recovery was defined as the ability of a fly to stand upright following paralysis and seizure.

### 2.3. Analysis of dendritic morphology in larval C4da neurons

Parental flies were maintained in standard culture vials and transferred to fresh media every 24 hours to prevent larval overcrowding and premature wandering. After approximately six days, motile wandering third-instar larvae that had not yet everted their anterior spiracles were selected for analysis. The *ppk-CD4-tdGFP* transgene (Han et al., 2011), derived from enhancer elements of the *Drosophila* DEG/ENaC gene *pickpocket1* (*ppk1*) (Adams et al., 1998), drives expression of membrane-localized GFP in class IV multiple-dendritic arborization (C4da) sensory neurons. Female larvae of the designated genotypes carrying *ppk-CD4-tdGFP* (e.g., *para^Shu^*/+; *ppk-CD4-tdGFP*/+) were dissected and filleted to remove internal tissues, then fixed in 4% paraformaldehyde in 1× PBS for 60 minutes at room temperature. Fixed body walls were incubated overnight at 4°C with an anti-GFP monoclonal antibody (12A6, Developmental Studies Hybridoma Bank, University of Iowa, Iowa City, IA) diluted in 1× PBT (1× PBS with 0.25% Triton X-100). Following four 10-minute washes in 1× PBT, samples were incubated overnight at 4°C with a secondary antibody (Alexa Fluor 488 goat anti-mouse, A11001, Invitrogen, Carlsbad, CA). After extensive washing in 1× PBT, body walls were mounted in Vectashield (Vector Labs, Newark, CA). Confocal images were acquired using a Zeiss 510 confocal microscope, generating 50–60 µm Z-stacks (5 µm per slice) that encompassed the entire dendritic arbor of a single dorsal ddaC C4da neuron. Maximum intensity projections of these Z-stacks were used as templates for dendritic tracing with the Simple Neurite Tracer plugin in Fiji (Schindelin et al., 2012). Traced images were analyzed using Sholl analysis (Sholl, 1953) and skeletonized to quantify branch and junction numbers in Fiji.

### 2.4. Analysis of sessile hemocytes in third instar larvae

Third-instar female larvae were prepared as described in section 2.3. Larvae were gently removed from the sides of culture vials using a damp paintbrush and transferred to a small Petri dish containing distilled water, where they were sexed, scored for FM7 with the actGFP balancer, and rinsed. To dry, larvae were placed on a small piece of KimWipe (Kimberly-Clark Professional, Roswell, GA). Once dried, they were transferred into a cell strainer, which was positioned inside a glass staining jar containing cotton balls soaked with 3 mL of isoflurane. Larvae were exposed to isoflurane vapor for 15 minutes before removal. Sessile hemocytes labeled with *He-GAL4* and *UAS-RedStinger* were imaged in intact larvae mounted on a glass slide between two stacks of three 18 × 18 mm cover glasses. The cover glass stacks were glued together and affixed to the slide with superglue, leaving a ∼0.25 cm gap between them. To prepare the imaging chamber, 90% glycerol in 1× PBS was added. A single larva was transferred from the cell strainer to the chamber using an inoculation loop moistened with glycerol mounting medium. The larva was positioned laterally (side-up), and a cover glass was placed on top and secured with painter’s tape. The chamber depth was adjusted so that the top cover glass just touched the larval cuticle. Segment A6 of each larva was imaged using a Zeiss 510 confocal microscope. Sessile hemocytes within the dendritic field of A6 C4da neurons were quantified using the ImageJ manual cell counter plugin.

### 2.5. Statistical Analysis

All data presented in this study were obtained from independent sets of experiments. “N” indicates the number of larvae or adult flies analyzed. Statistical analyses were conducted using GraphPad Prism 10 (GraphPad Software, Inc., Boston, MA). Fisher’s exact test with Bonferroni correction for multiple comparisons was applied to evaluate the effects of genetic manipulations on the downturned wing phenotype of paraShu mutants. Heat-induced seizure behavior was analyzed using two-way repeated-measures ANOVA, followed by Tukey’s multiple comparisons test as previously described (Lansdon et al., 2024). The severity of the bang-sensitive phenotype, dendritic morphology of C4da neurons, and the number of sessile hemocytes in the larval hematopoietic pocket were compared across genotypes using one-way ANOVA with Tukey’s multiple comparisons test.

## 3. Results

### 3.1. GstS1 knockdown in hemocytes suppresses hyperexcitable phenotypes in seizure-prone mutants, para^Shu^ and eas^2^

_T_o identify the specific tissues or cell types in which reduced *GstS1* function mitigates the hyperexcitable phenotypes of *para^Shu^*mutants, we examined the effects of *GstS1*-RNAi expression driven by tissue- or cell type–specific GAL4 drivers on their characteristic downturned wing phenotype. This wing posture abnormality in *para^Shu^* mutants is thought to arise from excessive muscle contraction secondary to neuronal hyperexcitability, as *Drosophila* muscles do not generate voltage-gated sodium currents (Kaas et al., 2016). Four GAL4 drivers were used: *nSyb-GAL4* for neurons, *repo-GAL4* for glia, *Lsp-GAL4* for fat bodies, and *He-GAL4* for hemocytes. Compared with the *GstS1-RNAi*–only control (*para^Shu^*/+; *UAS-GstS1-RNAi*/+, N = 112), the downturned wing phenotype in *para^Shu^* mutants was markedly suppressed when *GstS1-RNAi* expression was driven by *He-GAL4* (*para^Shu^*/+; *UAS-GstS1-RNAi*/*He-GAL4*, N = 61; Fisher’s exact test with Bonferroni correction for multiple comparisons, P_adj =_ 1.2 × 10^-40^) (Fig. 1A). This finding suggests the critical role of hemocytes in *GstS1*-mediated phenotypic suppression. In contrast, *GstS1* knockdown driven by other GAL4 drivers had no effect on the *para^Shu^* wing posture (N = 49, 98, and 48 for *repo-GAL4*, *nSyb-GAL4*, and *Lsp-GAL4*, respectively; P_adj >_ 0.05). Notably, the suppressive effect observed with *He-GAL4*–driven *GstS1-RNAi* was completely abolished by introducing a single copy of *srpHemo-GAL80*. Under these conditions, the wing phenotype reverted to the level observed in control *GstS1-RNAi*-only flies (N = 109; P_adj >_ 0.05) (Fig. 1A). Because *srpHemo-GAL80* antagonizes GAL4 activity specifically in hemocytes (Gyoergy et al., 2018), this result further confirmed a pivotal role for hemocytes in *GstS1*-mediated modification of *para^Shu^* wing posture phenotypes.

**Figure 1.**
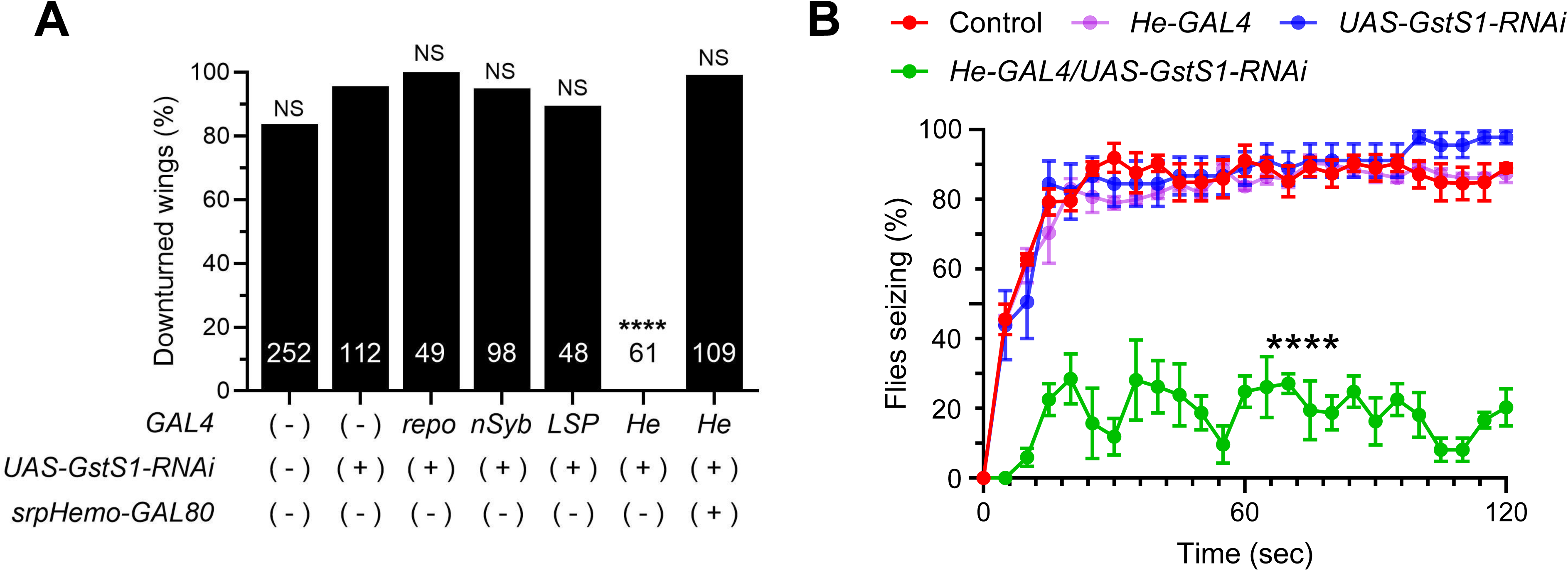
Knockdown of GstS1 in hemocytes suppresses the hyperexcitable phenotypes of adult para^Shu^ mutants. **(A)** Percentage of *para^Shu^* mutants exhibiting downturned wings when *GstS1-RNAi* is expressed in different cell types, compared with the RNAi-only control. The number of flies analyzed under each condition is shown within each bar. ****P < 0.0001; NS, not significant (P > 0.05). **(B)** Mean percentages (± SEM) of flies exhibiting seizures at each time point during exposure to 37°C. The numbers of flies tested for each condition were 74 (control, *para^Shu^*/+; +/+), 24 (*para^Shu^*/+; *GstS1-RNAi*/+), 114 (*para^Shu^*/+; *He-GAL4*/+), and 38 (*para^Shu^*/+; *GstS1-RNAi*/*He-GAL4*). ****P < 0.0001.

*para^Shu^* mutants show pronounced susceptibility to heat-induced seizures when exposed to elevated temperatures, 37°C (Kaas et al., 2016). After 2 minutes of exposure to 37°C, seizures occurred in 89.0%, 97.8%, and 87.4% of *para^Shu^* mutants (*para^Shu^*/+; +/+, N = 74), *para^Shu^*mutants with *GstS1-RNAi* only (*para^Shu^*/+; *UAS-GstS1-RNAi*/+, N = 24), and *para^Shu^* mutants with *He-GAL4* only (*para^Shu^*/+; *He-GAL4*/+, N = 114), respectively (Fig. 1B). In contrast, when *GstS1-RNAi* expression was driven by *He-GAL4* (*para^Shu^*/+; *UAS-GstS1-RNAi*/*He-GAL4*, N = 38), only 20.3% of flies exhibited seizures under the same condition (Fig. 1B). Compared with the other groups, *GstS1* knockdown in hemocytes had a statistically significant effect on heat-induced seizure behavior during the 2-min observation period (two-way repeated-measures ANOVA with Tukey’s multiple comparisons test, P < 0.0001), further underscoring the critical role of hemocyte-expressed *GstS1* in regulating seizure severity.

Suppression of hyperexcitability by hemocyte-specific *GstS1* knockdown was not limited to *para^Shu^* mutants. *eas^2^* is a loss-of-function allele of the X-linked ethanolamine kinase gene *easily shocked* (*eas*) (Pavlidis et al., 1994). When subjected to intense mechanical agitation, adult flies homozygous or hemizygous for the *eas^2^* mutation exhibit a characteristic bang-sensitive phenotype. This sequence of behaviors following a mechanical shock consists of initial uncoordinated movements, followed by transient paralysis, subsequent seizure activity, and eventual recovery (Pavlidis et al., 1994). The severity of the bang-sensitive phenotype can be quantified by measuring the time required for recovery from paralysis and seizure induced by mechanical stimulation (Chen et al., 2020; Kasuya et al., 2023).

We found that recovery time was shorter in *eas^2^* hemizygous mutant males (*eas^2^*/*Y*) carrying both *UAS-GstS1-RNAi* and *He-GAL4* (average ± SD, 76.7 ± 8.5 sec, N = 18) compared with those carrying only *UAS-GstS1-RNAi* (121.4 ± 26.6 sec, N = 19) or only *He-GAL4* (95.0 ± 15.7 sec, N = 19) (one-way ANOVA with Tukey’s multiple comparisons test, P_adj <_ 0.0001 and P_adj <_ 0.0148, respectively) (Fig. 2A). All *eas^2^* mutants carrying either *UAS-GstS1-RNAi* or *He-GAL4* alone (19 of 19 for each genotype) displayed vigorous wing beating prior to recovery, a typical feature of bang-sensitive mutants during this phase. In sharp contrast, none of the mutants carrying both *UAS-GstS1-RNAi* and *He-GAL4* (0 of 18) exhibited this hyperexcitable behavior (Fig. 2A, B), further demonstrating the seizure-suppressive effect of hemocyte-specific *GstS1* knockdown.

**Figure 2.**
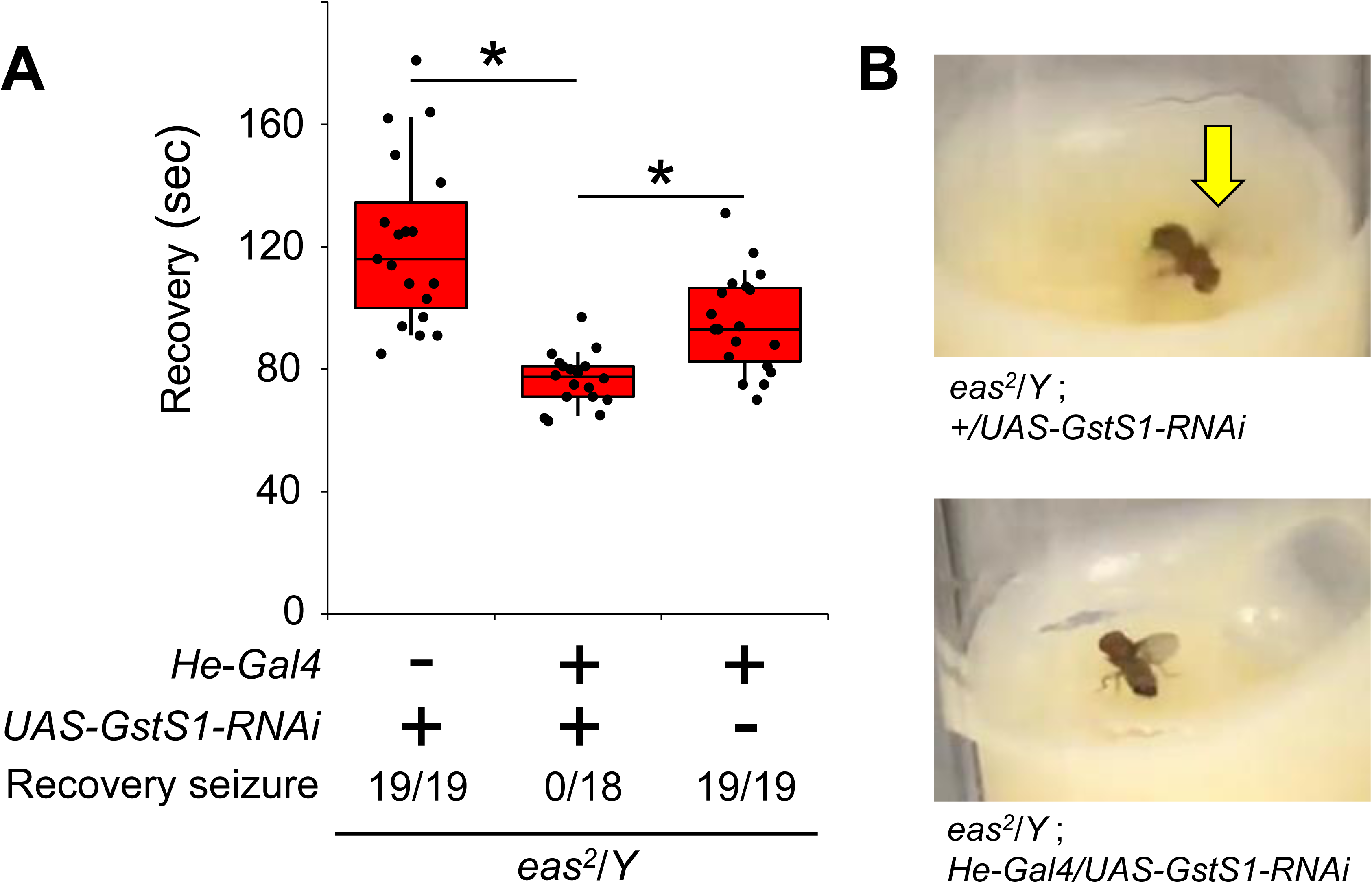
Knockdown of GstS1 in hemocytes reduces the severity of bang-sensitive phenotypes in adult eas mutants. **(A)** Mean recovery time (± SD) from mechanical shock–induced paralysis in hemizygous *eas^2^*/*Y* mutant males with or without *GstS1* knockdown in hemocytes. *P < 0.05; ****P < 0.0001. The numbers of flies tested under each condition, as well as those exhibiting recovery-associated convulsions, are indicated below each bar. **(B)** Representative images of *eas* mutant males during recovery from mechanical shock–induced paralysis with (top) and without (bottom) convulsive wing beating (red arrow).

### 3.2. Developmental knockdown of GstS1 effectively suppresses hyperexcitable phenotypes in adult para^Shu^ mutants

To determine when *GstS1* knockdown is required to suppress adult *para^Shu^* phenotypes, we conditionally reduced *GstS1* expression using *GstS1-RNAi* in combination with the GeneSwitch (GS)-GAL4 system. Activation of GAL4-mediated transcription by the *GS-GAL4* transgene requires the progesterone analog mifepristone (RU486) (Osterwalder et al., 2001) (Fig. 3A). When *GstS1* was ubiquitously knocked down only in adulthood—by raising *para^Shu^* mutants carrying both *tub-GS-GAL4* and *UAS-GstS1-RNAi* transgenes on control food during development and then switching them to RU486-containing food after eclosion—five-day-old *para^Shu^* adults continued to exhibit the downturned wing phenotype (V-R, 92.6%, N = 68). This frequency was comparable to that observed in *para^Shu^* mutants without *GstS1* knockdown (V-V, 95.0%, N = 60; Fisher’s exact test with Bonferroni correction, P_adj >_ 0.05). In contrast, *para^Shu^* mutants in which *GstS1* was knocked down only during development (R-V, 0%, N = 24) or during both development and adulthood (R-R, 0%, N = 15) did not display the downturned wing phenotype (Fig. 3B). The wing-posture phenotype of these groups (i.e., R-V and R-R) differed significantly from the control group (V-V), with P_adj v_alues of 1.4 × 10^-17^ and 1.1 × 10^-17^, respectively.

**Figure 3.**
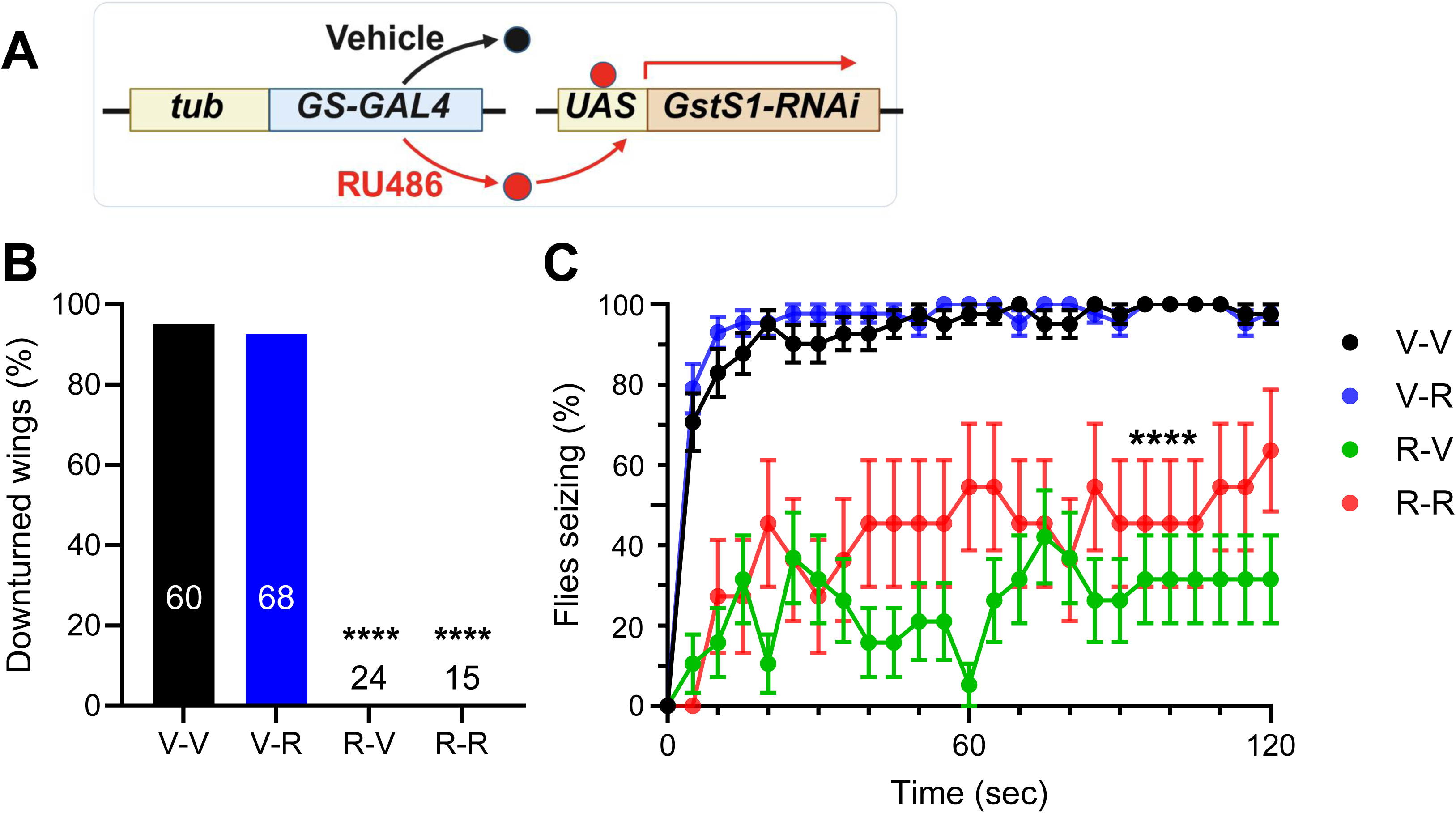
Knockdown of GstS1 during development suppresses hyperexcitability in adult para^Shu^ mutants. **(A)** Schematic diagram of the GeneSwitch (GS) system illustrating RU486-dependent induction of *GstS1-RNAi* expression driven by the tubulin enhancer. **(B)** Percentage of *para^Shu^* mutants exhibiting downturned wings when fed a vehicle (V) or RU486-supplemented (R) diet during larval development and/or adulthood. The numbers of flies tested under each condition are shown in parentheses. **(C)** Mean percentages (± SEM) of flies exhibiting seizures at each time point during exposure to 37°C. The numbers of flies analyzed for each condition were 41 (V-V, control), 43 (V-R), 19 (R-V), and 11 (R-R). ****P < 0.0001.

The effects of RU486-induced *GstS1* knockdown on heat-induced seizures were consistent with the results of the wing phenotype. When *GstS1* was knocked down only in adulthood (V-R, N = 43), seizure susceptibility was indistinguishable from controls (V-V, N = 41) (P > 0.05, two-way repeated measures ANOVA with Tukey’s multiple comparisons test). In contrast, seizure susceptibility was significantly reduced when *GstS1* was knocked down only during development (R-V, N = 19) or during both development and adulthood (R-R, N = 11) (P < 0.0001; Fig. 3C).

### 3.3. Knocking down GstS1 in hemocytes rescues abnormal dendrite growth in the larval sensory neurons of para^Shu^ mutants

The hyperexcitable phenotypes of adult *para^Shu^* mutants were suppressed when *GstS1* was knocked down during larval stages but not when it was knocked down during adulthood (Fig. 3). This finding suggests that *GstS1* function influences the behavioral phenotypes of adult mutants through its role in development of the nervous system. To explore the possible role of *GstS1* in neuronal development, we examined whether manipulation of *GstS1* affects the structure of larval class IV dendritic arborization (C4da) neurons in *para^Shu^* mutants. C4da neurons are a well-characterized class of sensory neurons composed of three bilaterally symmetrical cells—ddaC, v’ada, and vdab—per larval segment in the peripheral nervous system. They extend complex, non-overlapping dendritic arbors that tile the inner surface of the larval body wall and are widely used to study the developmental regulation of dendritic growth and patterning (Dong et al., 2015; Kanamori et al., 2015). Their stereotyped dendritic morphology can be reliably imaged and quantified, making them a suitable model for assessing how *GstS1* knockdown affects neuronal development in hyperexcitable mutants.

In our previous study, we demonstrated that C4da neurons in *para^Shu^*mutant larvae exhibit significantly greater dendritic complexity in regions proximal to the soma than those in WT larvae (Kasuya et al., 2023). To determine whether *GstS1* contributes to this phenotype, we visualized C4da neurons using the *ppk-CD4-tdGFP* transgene, which expresses a membrane-localized fluorescent marker specifically in these neurons (Han et al., 2011) (Fig. 4A). We then quantified dendritic architecture using Sholl analysis, which measures the number of dendritic intersections with concentric circles of increasing radius centered on the soma (Sholl, 1953). Quantification of the area under the Sholl curve (AUC) confirmed a significant increase in dendritic complexity within 10–100 μm of the soma in *para^Shu^* neurons (red) compared with WT neurons (white) (Fig. 4B and 4C). Consistent with this finding, analysis of reconstructed dendritic arbors showed that *para^Shu^* neurons possessed significantly more dendritic branches within 100 μm of the soma than WT neurons (Fig. 4D). Introducing a single copy of the *GstS1* null allele, *GstS1^M26^*, suppressed both phenotypes, restoring dendritic complexity and branch number to near-WT levels (yellow) (Fig. 4B, 4C and 4D). Similarly, hemocyte-specific knockdown of *GstS1* using the *He-GAL4* driver significantly reduced the dendritic complexity of *para^Shu^*neurons to WT levels (green) (Fig. 4B, 4C and 4D). Together, these findings indicate that *GstS1* activity in hemocytes promotes the increased dendritic complexity observed in *para^Shu^* C4da neurons, identifying hemocyte-derived *GstS1* as a key regulator of this neurodevelopmental phenotype.

**Figure 4.**
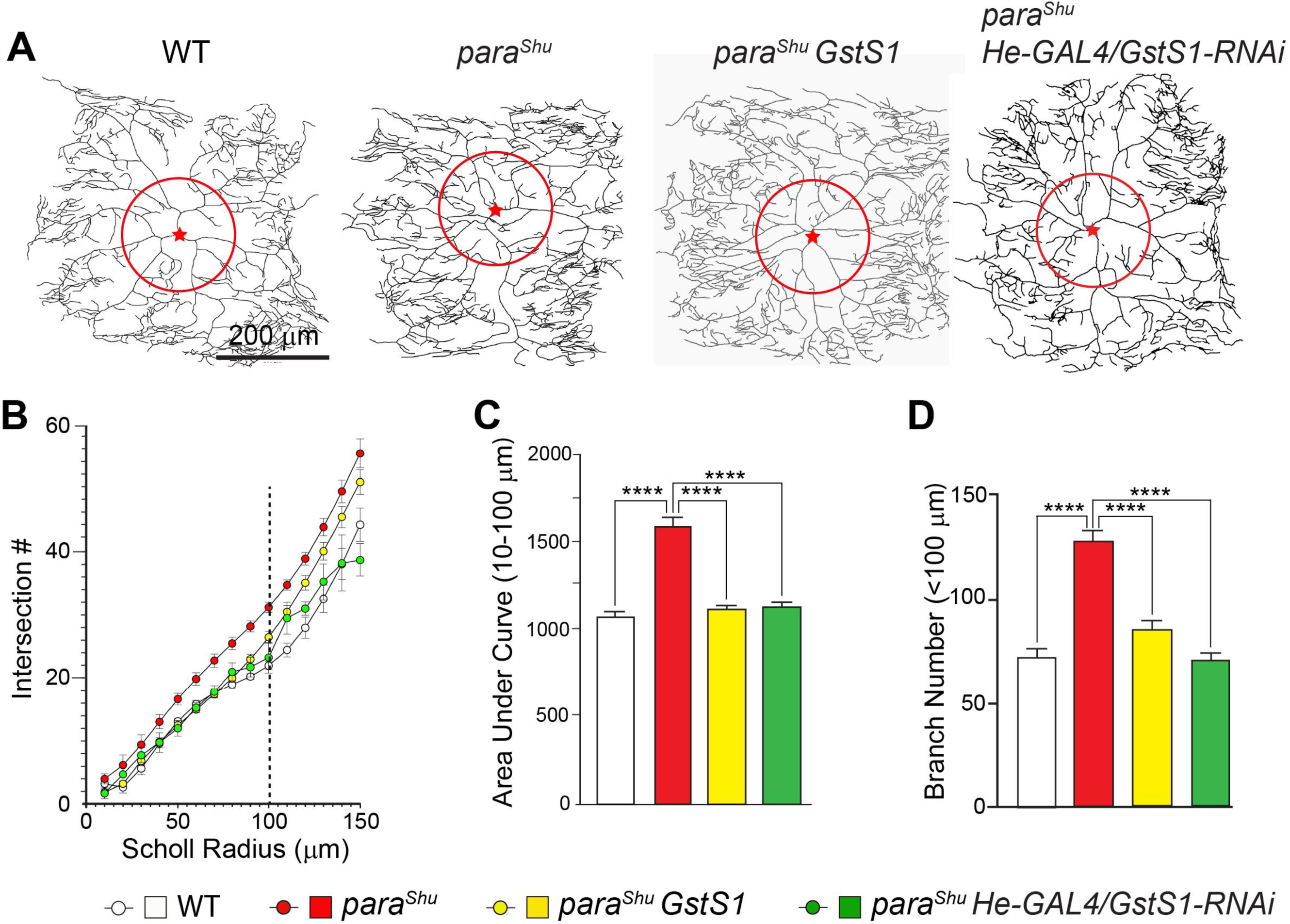
Increased dendritic complexity in para^Shu^ C4da neurons is suppressed by hemocyte-specific GstS1 knockdown. **(A)** Representative neurite traces of individual C4da neurons of indicated genotypes. The neuronal soma (red star) and a circle with a radius of 150 μm are shown for each neuron. **(B)** Scholl analysis indicating the number of dendritic branches intersecting circles with increasing radii from the soma Percentage of *para^Shu^* mutants exhibiting downturned wings when fed a vehicle (V) or RU486-supplemented (R) diet during larval development and/or adulthood. The numbers of flies tested under each condition are shown in parentheses. **(C)** Mean percentages (± SEM) of flies exhibiting seizures at each time point during exposure to 37°C. The numbers of flies analyzed for each condition were 41 (V-V, control), 43 (V-R), 19 (R-V), and 11 (R-R). ****P < 0.0001. (A) Mean recovery time (± SD) from mechanical shock–induced paralysis in hemizygous eas2/Y mutant males with or without GstS1 knockdown in hemocytes. *P < 0.05; ****P < 0.0001. The numbers of flies tested under each condition, as well as those exhibiting recovery-associated convulsions, are indicated below each bar. (B) Representative images of eas mutant males during recovery from mechanical shock–induced paralysis with (top) and without (bottom) convulsive wing beating (red arrow). (A) Percentage of paraShu mutants exhibiting downturned wings when GstS1-RNAi is expressed in different cell types, compared with the RNAi-only control. The number of flies analyzed under each condition is shown within each bar. ****P < 0.0001; NS, not significant (P > 0.05). (B) Mean percentages (± SEM) of flies exhibiting seizures at each time point during exposure to 37°C. The numbers of flies tested for each condition were 74 (control, paraShu/+; +/+), 24 (paraShu/+; GstS1-RNAi/+), 114 (paraShu/+; He-GAL4/+), and 38 (paraShu/+; GstS1-RNAi/He-GAL4). ****P < 0.0001.

### 3.4. The number of sessile hemocytes proximal to C4da neurons increases in para^Shu^ and eas^2^ mutants but is reduced by loss of GstS1 function

Given that hemocytes play a critical role in GstS1-mediated modulation of both adult seizure phenotypes (Fig. 1) and dendritic development of C4da neurons (Fig. 4), we next investigated whether and how hemocytes are influenced by seizure-causing mutations and reduced *GstS1* function. In *Drosophila* larvae, the hematopoietic pocket (HPP) is a segmentally repeated microenvironment situated between the body wall muscle and the epidermis, where macrophage-like sessile hemocytes reside and physically interact with sensory neurons, including C4da neurons (Makhijani et al., 2017; Makhijani et al., 2011).

To visualize and quantify hemocytes, we expressed the nuclear-targeted red fluorescent reporter Redstinger under the control of the hemocyte-specific *He-GAL4* driver. The number of hemocytes located within 150 μm of the C4da neuron soma (Fig. 5A) was quantified and compared across genotypes. We found that *para^Shu^*larvae exhibited a significant increase in hemocyte number near the C4da soma relative to WT controls (P < 0.0001, Fig. 5B). This phenotype was completely rescued either by introducing one copy of the *GstS1* null allele (*GstS1^M26^*) or by hemocyte-specific knockdown of *GstS1* using *He-GAL4* and *UAS-GstS1-RNAi* (Fig. 5B). A similar increase in sessile hemocytes near the C4da soma was observed in *eas^2^* mutants (P < 0.0001, Fig. 5C), and this phenotype was likewise suppressed by *GstS1^M26^*(Fig. 5C). These results indicate that hyperexcitable mutations such as *para^Shu^* and *eas^2^*alter hemocyte localization in a GstS1-dependent manner. Consistent with the parallel suppressive effects of reduced *GstS1* function and dietary ALA supplementation on the adult *para^Shu^* seizure phenotype (Chen et al., 2020; Kasuya et al., 2023), ALA feeding also decreased the number of sessile hemocytes in the HPP in a dose-dependent manner (Fig. 5D). Together, these findings suggest that modulation of hemocyte distribution represents a shared mechanism by which GstS1 activity and dietary factors influence neural excitability and associated phenotypes.

**Figure 5.**
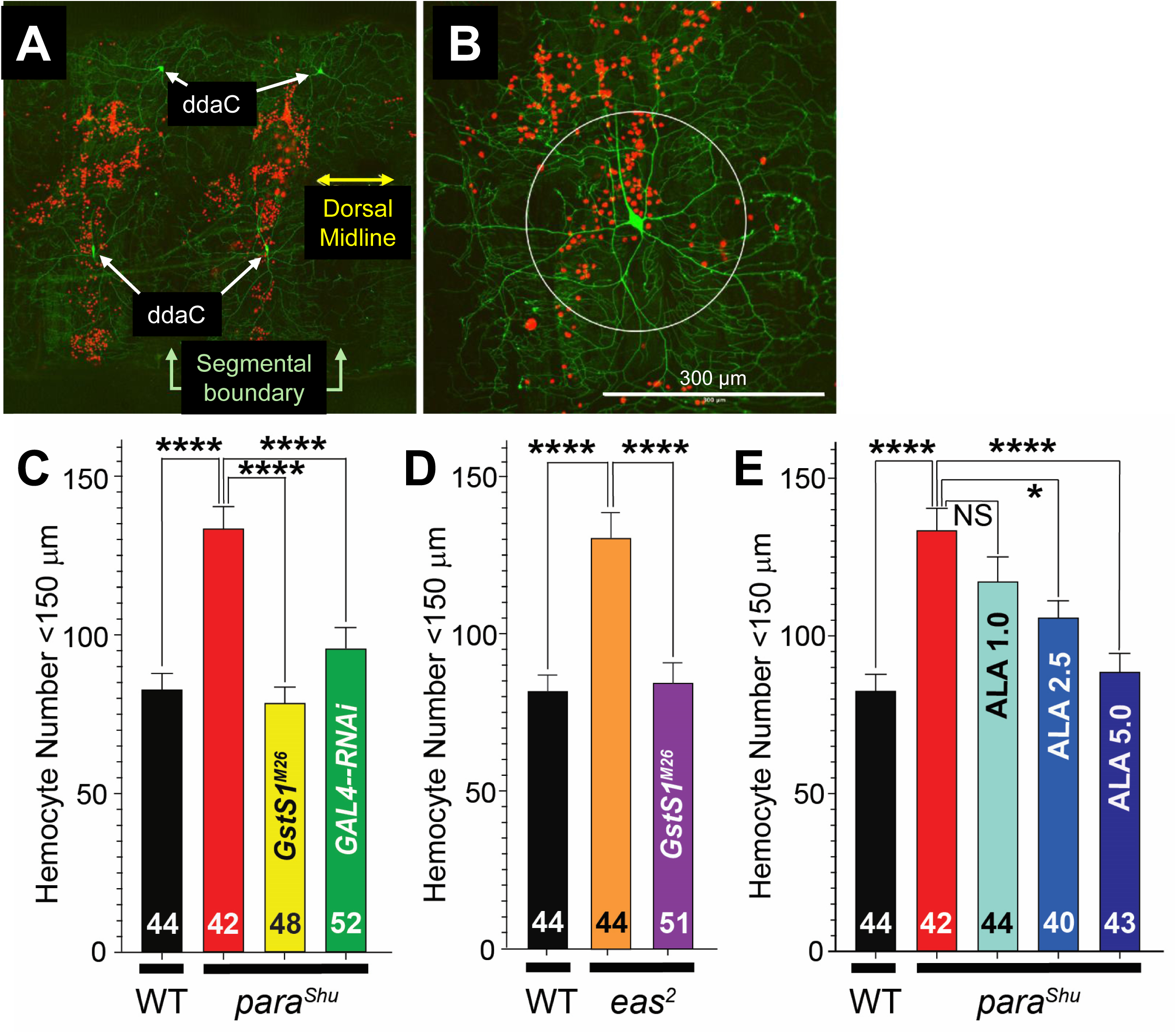

## 4. Discussion

A comprehensive understanding of how epileptic phenotypes arise—and how they are shaped by both genetic and environmental factors—is essential for developing effective strategies to prevent and treat epilepsy. Seizure-prone *Drosophila* mutants provide a powerful experimental platform for identifying and characterizing modifiers of neuronal hyperexcitability, largely due to precisely controlled genetic backgrounds and environmental conditions (Chen et al., 2020; Hekmat-Scafe et al., 2005; Kasuya et al., 2019; Kasuya et al., 2023; Lansdon et al., 2024; Lin et al., 2017; Lin et al., 2018; Mee et al., 2004; Song et al., 2008). Building on our previous finding that a global reduction in *GstS1* function suppresses hyperexcitable phenotypes in seizure-prone *Drosophila* mutants (Chen et al., 2020), the present study represents an important step toward defining the cellular and developmental mechanisms underlying this effect.

By systematically reducing *GstS1* function in specific cell types, we identified hemocytes—not neurons or glia—as the critical site mediating GstS1-dependent suppression of seizure phenotypes in both *para^Shu^*and *eas^2^* mutants. *Drosophila* hemocytes are well known for their roles in nervous system development, including the regulation of synaptic pruning, extracellular matrix remodeling, and the secretion of trophic factors (Matsubayashi et al., 2017; Morin-Poulard et al., 2021; Sung et al., 2024). Our findings extend these roles by demonstrating that GstS1 activity in hemocytes influences neuronal excitability later in life. Together, these results highlight immune cells as potent modulators of neural circuit development and function.

The mechanism by which GstS1 acts in hemocytes likely involves the regulation of bioactive lipid mediators. GstS1 is considered an ortholog of mammalian hematopoietic prostaglandin D synthase (HPGDS) (Scarpati et al., 2019), an enzyme that produces prostaglandin D2 (PGD2), a lipid mediator implicated in inflammation, oxidative stress, and neuronal survival (Liu et al., 2008; Ricciotti & FitzGerald, 2011; Taniguchi et al., 2007). Comparable functions have been described in other insects; for example, knockdown of the *GstS1* counterpart in the lepidopteran insect *Spodoptera exigua* (Spodoptera prostaglandin D synthase, SePGDS) disrupts hemocyte function and immune responses, defects that can be rescued by PGD2 supplementation (Sajjadian et al., 2020). In *Drosophila*, PGD2 production has been detected when arachidonic acid is supplied (Fujinaga et al., 2025), further supporting the notion that GstS1 participates in prostaglandin-related lipid signaling. Although the precise physiological roles of PGD2 in flies remain to be fully defined, our data support a model in which hemocyte-derived GstS1 regulates lipid signaling that modulates neuronal excitability through paracrine or humoral mechanisms.

Several prior observations support this interpretation. *GstS1* overexpression in hemocytes increases hemocyte abundance (Stofanko et al., 2008), and *GstS1* expression in hemocytes is strongly upregulated during metamorphosis (Regan et al., 2013). Moreover, we previously showed that innate immune genes, including antimicrobial peptide genes, are upregulated in *para^Shu^* mutants, and that reduced *GstS1* function reverses this immune activation (Chen et al., 2020; Kaas et al., 2016). Given the well-established link between immune activity and seizure susceptibility across species (Nukala et al., 2023; Annamaria Vezzani et al., 2019; A. Vezzani et al., 2011), these findings further support a role for GstS1 in regulating neuronal hyperexcitability through neuroimmune interactions. By demonstrating that macrophage-like hemocytes influence neuronal excitability, this study establishes *Drosophila* as a genetically tractable model for dissecting immune–metabolic contributions to hyperexcitable states.

Our temporal manipulation experiments showed that conditional *GstS1* knockdown before adult eclosion—but not afterward—was sufficient to suppress *para^Shu^* adult phenotypes, indicating that *GstS1* function is required during neural circuit formation. Hemocytes are abundant during the larval and pupal stages, when they patrol developing tissues, clear apoptotic debris, and maintain basement membrane integrity (Gold & Brückner, 2015; Lemaitre & Hoffmann, 2007). Altered GstS1 activity during this developmental window may shift hemocyte lipid metabolism toward a less inflammatory or more neuroprotective state, thereby permitting more normal neural wiring in seizure-prone mutants. Once neural circuits are established, however, reducing GstS1 activity may no longer reverse these developmental effects. This developmental specificity parallels mammalian studies showing that early-life immune perturbations increase seizure susceptibility later in life (Auvin et al., 2010; Rana & Musto, 2018; Rojas et al., 2019).

Consistent with this model, hemocyte-specific *GstS1* knockdown rescued dendritic overgrowth in larval C4da neurons of *para^Shu^* mutants, linking neuroimmune signaling to structural plasticity during development. C4da neurons normally exhibit highly stereotyped dendritic tiling governed by intrinsic transcriptional programs and extrinsic cues, including signals from epidermal and immune cells (Corty et al., 2009; Igaki & Miura, 2014). In hyperexcitable mutants, neuronal stress may disrupt these interactions, promoting aberrant dendritic growth. Reducing GstS1 activity in hemocytes appears to normalize these interactions, restoring proper dendritic patterning. Notably, seizure-prone mutants displayed increased numbers of sessile hemocytes near neuronal somata, suggesting that neuronal hyperexcitability may recruit hemocytes to neural surfaces in a manner analogous to immune surveillance in vertebrate systems.

Notably, the seizure-suppressive effects of hemocyte-specific *GstS1* knockdown closely resemble those of dietary ALA supplementation, including a shared developmental window of effectiveness (Chen et al., 2020; Kasuya et al., 2023). Both interventions also reduce hemocyte accumulation near neurons (Fig. 5). Given that ALA is an ω-3 polyunsaturated fatty acid (PUFA) and GstS1 metabolizes lipid substrates, a plausible explanation is that GstS1 regulates the balance between ω-6–and ω-3–derived lipid mediators. In mammals, ω-6 PUFAs generally give rise to pro-inflammatory prostaglandins, whereas ω-3 PUFAs generate anti-inflammatory and pro-resolving mediators (Basil & Levy, 2016; Ricciotti & FitzGerald, 2011). If GstS1 functions as a prostaglandin D synthase, reduced GstS1 activity may attenuate ω-6–derived pro-inflammatory signaling, thereby phenocopying ω-3 PUFA enrichment. Consistent with this model, metabolomic analyses show that ALA supplementation reverses oxidative stress–and immune-related metabolic abnormalities in *para^Shu^*mutants (Kruth et al., 2025). Together, these findings suggest that genetic and dietary interventions converge on a shared lipid-mediated pathway that maintains neuroimmune balance and limits hyperexcitability.

In summary, our results support a unified model in which GstS1 modulates hemocyte-derived lipid signals that shape nervous system excitability through developmental mechanisms. Reduced GstS1 activity may alter hemocyte lipid metabolism in a manner that promotes the formation of neural circuits that dampen physiological and behavioral hyperexcitability. Importantly, the finding that a lipid-metabolizing enzyme in immune cells modifies genetically induced hyperexcitability broadens our understanding of seizure modifiers. In human epilepsies caused by defective *SCN1A* and related channelopathies, substantial clinical variability among patients carrying identical mutations suggests that immune–metabolic interactions may function as a modifier axis (Martins Custodio et al., 2023; Annamaria Vezzani et al., 2019; Zuberi et al., 2011). Thus, the *Drosophila* model provides a valuable system for gaining fundamental insight into how lipid-mediated immune signaling contributes to the regulation of neural development and nervous system excitability.

Although our results establish a strong connection between GstS1, hemocytes, and neural excitability, several important questions remain. In particular, the specific biochemical products generated by GstS1 in hemocytes that mediate phenotypic suppression have not yet been identified, and lipidomic analyses will be essential to define prostaglandin-like or other lipid metabolites altered by *GstS1* knockdown. It also remains unclear whether hemocytes influence neurons primarily through direct cell–cell interactions or indirectly via diffusible signaling molecules. Approaches combining live imaging of hemocyte–neuron interactions with single-cell transcriptomic analyses should help resolve these mechanisms and define hemocyte activation states associated with nervous system hyperexcitability. In addition, while C4da neurons provide a useful developmental readout, it will be important to determine whether similar hemocyte–neuron interactions occur within central neural circuits that directly govern adult seizure behaviors. Finally, elucidating how GstS1 interfaces with canonical immune signaling pathways, including Toll, Imd, and JAK–STAT, will clarify whether *GstS1*- and ALA-dependent effects act upstream of, downstream of, or in parallel with established immune programs.

## References

Adams, C. M., Anderson, M. G., Motto, D. G., Price, M. P., Johnson, W. A., & Welsh, M. J. (1998). Ripped pocket and pickpocket, novel Drosophila DEG/ENaC subunits expressed in early development and in mechanosensory neurons. J Cell Biol, 140(1), 143–152. 10.1083/jcb.140.1.143

Auvin, S., Shin, D., Mazarati, A., & Sankar, R. (2010). Inflammation induced by LPS enhances epileptogenesis in immature rat and may be partially reversed by IL1RA. Epilepsia, 51 Suppl 3(Suppl 3), 34–38. 10.1111/j.1528-1167.2010.02606.x

Basil, M. C., & Levy, B. D. (2016). Specialized pro-resolving mediators: endogenous regulators of infection and inflammation. Nat Rev Immunol, 16(1), 51–67. 10.1038/nri.2015.4

Calder, P. C. (2017). Omega-3 fatty acids and inflammatory processes: from molecules to man. Biochem Soc Trans, 45(5), 1105–1115. 10.1042/bst20160474

Chen, H. L., Kasuya, J., Lansdon, P., Kaas, G., Tang, H., Sodders, M., & Kitamoto, T. (2020). Reduced Function of the Glutathione S-Transferase S1 Suppresses Behavioral Hyperexcitability in Drosophila Expressing Mutant Voltage-Gated Sodium Channels. G3 (Bethesda), 10(4), 1327–1340. 10.1534/g3.119.401025

Corty, M. M., Matthews, B. J., & Grueber, W. B. (2009). Molecules and mechanisms of dendrite development in Drosophila. Development, 136(7), 1049–1061. 10.1242/dev.014423

Dong, X., Shen, K., & Bulow, H. E. (2015). Intrinsic and extrinsic mechanisms of dendritic morphogenesis. Annu Rev Physiol, 77, 271–300. 10.1146/annurev-physiol-021014-071746

Fischer, F. P., Karge, R. A., Weber, Y. G., Koch, H., Wolking, S., & Voigt, A. (2023). Drosophila melanogaster as a versatile model organism to study genetic epilepsies: An overview. Front Mol Neurosci, 16, 1116000. 10.3389/fnmol.2023.1116000

Fujinaga, D., Nolan, C., & Yamanaka, N. (2025). Functional characterization of eicosanoid signaling in Drosophila development. PLoS Genetics, 21(5), e1011705. 10.1371/journal.pgen.1011705

Gold, K. S., & Brückner, K. (2015). Macrophages and cellular immunity in Drosophila melanogaster. Seminars in Immunology, 27(6), 357–368. 10.1016/j.smim.2016.03.010

Gyoergy, A., Roblek, M., Ratheesh, A., Valoskova, K., Belyaeva, V., Wachner, S., Matsubayashi, Y., Sanchez-Sanchez, B. J., Stramer, B., & Siekhaus, D. E. (2018). Tools Allowing Independent Visualization and Genetic Manipulation of Drosophila melanogaster Macrophages and Surrounding Tissues. G3 (Bethesda), 8(3), 845–857. 10.1534/g3.117.300452

Han, C., Jan, L. Y., & Jan, Y. N. (2011). Enhancer-driven membrane markers for analysis of nonautonomous mechanisms reveal neuron-glia interactions in Drosophila. Proc Natl Acad Sci U S A, 108(23), 9673–9678. 10.1073/pnas.1106386108 1106386108 [pii]

Hayes, J. D., Flanagan, J. U., & Jowsey, I. R. (2005). Glutathione transferases. Annu Rev Pharmacol Toxicol, 45, 51–88. 10.1146/annurev.pharmtox.45.120403.095857

Hekmat-Scafe, D. S., Dang, K. N., & Tanouye, M. A. (2005). Seizure suppression by gain-of-function escargot mutations. Genetics, 169(3), 1477–1493. http://www.ncbi.nlm.nih.gov/entrez/query.fcgi?cmd=Retrieve&db=PubMed&dopt=Citation&list_uids=15654097

Igaki, T., & Miura, M. (2014). The Drosophila TNF ortholog Eiger: emerging physiological roles and evolution of the TNF system. Semin Immunol, 26(3), 267–274. 10.1016/j.smim.2014.05.003

Kaas, G. A., Kasuya, J., Lansdon, P., Ueda, A., Iyengar, A., Wu, C. F., & Kitamoto, T. (2016). Lithium-Responsive Seizure-Like Hyperexcitability Is Caused by a Mutation in the Drosophila Voltage-Gated Sodium Channel Gene paralytic. eNeuro, 3(5). 10.1523/ENEURO.0221-16.2016

Kanamori, T., Togashi, K., Koizumi, H., & Emoto, K. (2015). Dendritic Remodeling: Lessons from Invertebrate Model Systems. Int Rev Cell Mol Biol, 318, 1–25. 10.1016/bs.ircmb.2015.05.001

Kasuya, J., Iyengar, A., Chen, H. L., Lansdon, P., Wu, C. F., & Kitamoto, T. (2019). Milk-whey diet substantially suppresses seizure-like phenotypes of para(Shu), a Drosophila voltage-gated sodium channel mutant. J Neurogenet, 33(3), 164–178. 10.1080/01677063.2019.1597082

Kasuya, J., Johnson, W., Chen, H. L., & Kitamoto, T. (2023). Dietary Supplementation with Milk Lipids Leads to Suppression of Developmental and Behavioral Phenotypes of Hyperexcitable Drosophila Mutants. Neuroscience, 520, 1–17. 10.1016/j.neuroscience.2023.03.027

Klassen, T., Davis, C., Goldman, A., Burgess, D., Chen, T., Wheeler, D., McPherson, J., Bourquin, T., Lewis, L., Villasana, D., Morgan, M., Muzny, D., Gibbs, R., & Noebels, J. (2011). Exome sequencing of ion channel genes reveals complex profiles confounding personal risk assessment in epilepsy. Cell, 145(7), 1036–1048. 10.1016/j.cell.2011.05.025

Kluckova, D., Kolnikova, M., Lacinova, L., Jurkovicova-Tarabova, B., Foltan, T., Demko, V., Kadasi, L., Ficek, A., & Soltysova, A. (2020). A Study among the Genotype, Functional Alternations, and Phenotype of 9 SCN1A Mutations in Epilepsy Patients. Sci Rep, 10(1), 10288. 10.1038/s41598-020-67215-y

Kruth, K., Kasuya, J., Hand, V., Iyengar, A., & Kitamoto, T. (2025). Effects of a Gain-of-Function Mutation in the Voltage-Gated Sodium Channel Gene, and of Dietary α-Linolenic Acid Supplementation, on Whole-Body Metabolism in Drosophila. Res Sq. 10.21203/rs.3.rs-7786194/v1

Lansdon, P., Kasuya, J., & Kitamoto, T. (2024). Commensal bacteria exacerbate seizure-like phenotypes in Drosophila voltage-gated sodium channel mutants. Genes Brain Behav, 23(5), e70000. 10.1111/gbb.70000

Lemaitre, B., & Hoffmann, J. (2007). The host defense of Drosophila melanogaster. Annu Rev Immunol, 25, 697–743. http://www.ncbi.nlm.nih.gov/entrez/query.fcgi?cmd=Retrieve&db=PubMed&dopt=Citation&list_uids=17201680

Lewis, E. B. (1960). A new standard food medium. Drosophila Information Service, 34, 117–118.

Lin, W. H., Giachello, C. N., & Baines, R. A. (2017). Seizure control through genetic and pharmacological manipulation of Pumilio in Drosophila: a key component of neuronal homeostasis. Dis Model Mech, 10(2), 141–150. 10.1242/dmm.027045

Lin, W. H., He, M., Fan, Y. N., & Baines, R. A. (2018). An RNAi-mediated screen identifies novel targets for next-generation antiepileptic drugs based on increased expression of the homeostatic regulator pumilio. J Neurogenet, 32(2), 106–117. 10.1080/01677063.2018.1465570

Liu, H., Dinkova-Kostova, A. T., & Talalay, P. (2008). Coordinate regulation of enzyme markers for inflammation and for protection against oxidants and electrophiles. Proc Natl Acad Sci U S A, 105(41), 15926–15931. 10.1073/pnas.0808346105

Makhijani, K., Alexander, B., Rao, D., Petraki, S., Herboso, L., Kukar, K., Batool, I., Wachner, S., Gold, K. S., Wong, C., O’Connor, M. B., & Bruckner, K. (2017). Regulation of Drosophila hematopoietic sites by Activin-beta from active sensory neurons. Nat Commun, 8, 15990. 10.1038/ncomms15990

Makhijani, K., Alexander, B., Tanaka, T., Rulifson, E., & Bruckner, K. (2011). The peripheral nervous system supports blood cell homing and survival in the Drosophila larva. Development, 138(24), 5379–5391. 10.1242/dev.067322 dev.067322 [pii]

Martins Custodio, H., Clayton, L. M., Bellampalli, R., Pagni, S., Silvennoinen, K., Caswell, R., Consortium, G. E. R., Brunklaus, A., Guerrini, R., Koeleman, B. P. C., Lemke, J. R., Møller, R. S., Scheffer, I. E., Weckhuysen, S., Zara, F., Zuberi, S., Kuchenbaecker, K., Balestrini, S., Mills, J. D., & Sisodiya, S. M. (2023). Widespread genomic influences on phenotype in Dravet syndrome, a ‘monogenic’ condition. Brain, 146(9), 3885–3897. 10.1093/brain/awad111

Matsubayashi, Y., Louani, A., Dragu, A., Sánchez-Sánchez, B. J., Serna-Morales, E., Yolland, L., Gyoergy, A., Vizcay, G., Fleck, R. A., Heddleston, J. M., Chew, T.-L., Siekhaus, D. E., & Stramer, B. M. (2017). A Moving Source of Matrix Components Is Essential for De Novo Basement Membrane Formation. Current Biology, 27(22), 3526–3534.e3524. 10.1016/j.cub.2017.10.001

Mee, C. J., Pym, E. C., Moffat, K. G., & Baines, R. A. (2004). Regulation of neuronal excitability through pumilio-dependent control of a sodium channel gene. J Neurosci, 24(40), 8695–8703. 10.1523/JNEUROSCI.2282-04.2004

Morin-Poulard, I., Tian, Y., Vanzo, N., & Crozatier, M. (2021). Drosophila as a Model to Study Cellular Communication Between the Hematopoietic Niche and Blood Progenitors Under Homeostatic Conditions and in Response to an Immune Stress. Front Immunol, 12, 719349. 10.3389/fimmu.2021.719349

Ng, A. C., Chahine, M., Scantlebury, M. H., & Appendino, J. P. (2024). Channelopathies in epilepsy: an overview of clinical presentations, pathogenic mechanisms, and therapeutic insights. J Neurol, 271(6), 3063–3094. 10.1007/s00415-024-12352-x

Nukala, K. M., Lilienthal, A. J., Lye, S. H., Bassuk, A. G., Chtarbanova, S., & Manak, J. R. (2023). Downregulation of oxidative stress-mediated glial innate immune response suppresses seizures in a fly epilepsy model. Cell Rep, 42(1), 112004. 10.1016/j.celrep.2023.112004

Osterwalder, T., Yoon, K. S., White, B. H., & Keshishian, H. (2001). A conditional tissue-specific transgene expression system using inducible GAL4. Proceedings of the National Academy of Sciences, 98(22), 12596–12601. 10.1073/pnas.221303298

Pavlidis, P., Ramaswami, M., & Tanouye, M. A. (1994). The Drosophila easily shocked gene: a mutation in a phospholipid synthetic pathway causes seizure, neuronal failure, and paralysis. Cell, 79(1), 23–33. http://www.ncbi.nlm.nih.gov/pubmed/7923374

Rana, A., & Musto, A. E. (2018). The role of inflammation in the development of epilepsy. Journal of Neuroinflammation, 15(1), 144. 10.1186/s12974-018-1192-7

Regan, J. C., Brandao, A. S., Leitao, A. B., Mantas Dias, A. R., Sucena, E., Jacinto, A., & Zaidman-Remy, A. (2013). Steroid hormone signaling is essential to regulate innate immune cells and fight bacterial infection in Drosophila. PLoS Pathog, 9(10), e1003720. 10.1371/journal.ppat.1003720

Ricciotti, E., & FitzGerald, G. A. (2011). Prostaglandins and inflammation. Arterioscler Thromb Vasc Biol, 31(5), 986–1000. 10.1161/ATVBAHA.110.207449

Rojas, A., Chen, D., Ganesh, T., Varvel, N. H., & Dingledine, R. (2019). The COX-2/prostanoid signaling cascades in seizure disorders. Expert Opinion on Therapeutic Targets, 23(1), 1–13. 10.1080/14728222.2019.1554056

Sajjadian, S. M., Ahmed, S., Al Baki, M. A., & Kim, Y. (2020). Prostaglandin D(2) synthase and its functional association with immune and reproductive processes in a lepidopteran insect, Spodoptera exigua. Gen Comp Endocrinol, 287, 113352. 10.1016/j.ygcen.2019.113352

Salinas, A. E., & Wong, M. G. (1999). Glutathione S-transferases--a review. Curr Med Chem, 6(4), 279–309.

Scarpati, M., Qi, Y., Govind, S., & Singh, S. (2019). A combined computational strategy of sequence and structural analysis predicts the existence of a functional eicosanoid pathway in Drosophila melanogaster. PLoS One, 14(2), e0211897. 10.1371/journal.pone.0211897

Schindelin, J., Arganda-Carreras, I., Frise, E., Kaynig, V., Longair, M., Pietzsch, T., Preibisch, S., Rueden, C., Saalfeld, S., Schmid, B., Tinevez, J. Y., White, D. J., Hartenstein, V., Eliceiri, K., Tomancak, P., & Cardona, A. (2012). Fiji: an open-source platform for biological-image analysis. Nat Methods, 9(7), 676–682. 10.1038/nmeth.2019

Sholl, D. A. (1953). Dendritic organization in the neurons of the visual and motor cortices of the cat. J Anat, 87(4), 387–406. https://www.ncbi.nlm.nih.gov/pubmed/13117757

Song, J., Parker, L., Hormozi, L., & Tanouye, M. A. (2008). DNA topoisomerase I inhibitors ameliorate seizure-like behaviors and paralysis in a Drosophila model of epilepsy. Neuroscience, 156(3), 722–728. 10.1016/j.neuroscience.2008.07.024

Stofanko, M., Kwon, S. Y., & Badenhorst, P. (2008). A misexpression screen to identify regulators of Drosophila larval hemocyte development. Genetics, 180(1), 253–267. 10.1534/genetics.108.089094

Sun, L., Gilligan, J., Staber, C., Schutte, R. J., Nguyen, V., O’Dowd, D. K., & Reenan, R. (2012). A knock-in model of human epilepsy in Drosophila reveals a novel cellular mechanism associated with heat-induced seizure. J Neurosci, 32(41), 14145–14155. 10.1523/JNEUROSCI.2932-12.2012 32/41/14145 [pii]

Sung, H. H., Li, H., Huang, Y. C., Ai, C. L., Hsieh, M. Y., Jan, H. M., Peng, Y. J., Lin, H. Y., Yeh, C. H., Lin, S. Y., Yeh, C. Y., Cheng, Y. J., Khoo, K. H., Lin, C. H., & Chien, C. T. (2024). Galectins induced from hemocytes bridge phosphatidylserine and N-glycosylated Drpr/CED-1 receptor during dendrite pruning. Nat Commun, 15(1), 7402. 10.1038/s41467-024-51581-6

Taniguchi, H., Mohri, I., Okabe-Arahori, H., Aritake, K., Wada, K., Kanekiyo, T., Narumiya, S., Nakayama, M., Ozono, K., Urade, Y., & Taniike, M. (2007). Prostaglandin D2 protects neonatal mouse brain from hypoxic ischemic injury. J Neurosci, 27(16), 4303–4312. 10.1523/JNEUROSCI.0321-07.2007

Vaish, S., Gupta, D., Mehrotra, R., Mehrotra, S., & Basantani, M. K. (2020). Glutathione S-transferase: a versatile protein family. 3 Biotech, 10(7), 321. 10.1007/s13205-020-02312-3

Vezzani, A., Balosso, S., & Ravizza, T. (2019). Neuroinflammatory pathways as treatment targets and biomarkers in epilepsy. Nature Reviews Neurology, 15(8), 459–472. 10.1038/s41582-019-0217-x

Vezzani, A., French, J., Bartfai, T., & Baram, T. Z. (2011). The role of inflammation in epilepsy. Nat Rev Neurol, 7(1), 31–40. 10.1038/nrneurol.2010.178

Williamson, R. L. (1982). Lithium stops hereditary shuddering in Drosophila melanogaster. Psychopharmacology (Berl), 76(3), 265–268. http://www.ncbi.nlm.nih.gov/entrez/query.fcgi?cmd=Retrieve&db=PubMed&dopt=Citation&list_uids=6808547

Zhang, H., Tan, J., Reynolds, E., Kuebler, D., Faulhaber, S., & Tanouye, M. (2002). The Drosophila slamdance gene: a mutation in an aminopeptidase can cause seizure, paralysis and neuronal failure. Genetics, 162(3), 1283–1299. http://www.ncbi.nlm.nih.gov/pubmed/12454073

Zuberi, S. M., Brunklaus, A., Birch, R., Reavey, E., Duncan, J., & Forbes, G. H. (2011). Genotype-phenotype associations in SCN1A-related epilepsies. Neurology, 76(7), 594–600. 10.1212/WNL.0b013e31820c309b

